# De novo design of ATPase based on the blueprint optimized for harboring the P-loop motif

**DOI:** 10.1101/2024.10.03.616451

**Authors:** Takahiro Kosugi, Mikio Tanabe, Nobuyasu Koga

## Abstract

De novo design of proteins has seen remarkable recent progress and has provided understanding of folding and functional expression. However, rationally creating enzymes with high activity comparable to most naturally occurring enzymes remains challenging. Here, we attempted to design an ATPase de novo, through the exploration of an optimal backbone blueprint to incorporate a conserved phosphate binding motif, the P-loop, into designed structures. The designed protein, based on the identified blueprint, was found to be a monomer with high thermal stability, and exhibited the ATPase ability. The crystal structure was closely matched to the design model, both at the overall structure level and within the P-loop motif. Interestingly, AlphaFold was not able to predict the designed structure accurately, indicating the difficulties of predicting folded structures for novel protein sequences. Remarkably, the designed protein exhibited ATPase ability even at temperatures around 100 °C, with significantly increased activity. However, the ATPase activity was still not comparable to those of naturally occurring enzymes. This suggests that the P-loop motif alone is insufficient to achieve the high ATPase activity seen in naturally occurring enzymes, indicating that other structural components are necessary to reach such activity levels.

## Introduction

De novo protein design provides opportunities to explore fundamental understanding of folding and functional expression programmed into protein structures. A variety of proteins have been designed from scratch, together with the understanding of principles for folding and functional expression (2; 9; 33; 3; 11). However, computationally creating highly active enzymes without further experimental optimizations such as directed evolution or site-directed mutagenesis remains an unsolved problem (18; 17). In this research, we aimed to uncover the design principles to create ATPase with high activity through de novo design, particularly focused on an optimal backbone blueprint, involving secondary structure lengths, loop patterns, and registries between β-strands, to incorporate a conserved phosphate-binding motif, known as the P-loop motif.

Many naturally occurring ATPases (30; 25) possess the P-loop motif, with a sequence pattern of GX_1_X_2_X_3_X_4_GK[T/S] (X indicates any kind of amino acid residues). Previously, Romero et al. designed proteins with ATPase activity (24). In their study, they discovered that inserting the conserved sequence into a loop region of a de novo designed protein scaffold is sufficient to provide ATP hydrolase ability. However, the ATPase activity of designed proteins, approximately one ATP molecule every 30 minutes, is significantly lower compared to that of naturally occurring enzymes (18), prompting further investigation into how to improve catalytic activity. Interestingly, the inserted P-loop sequence exhibits unstable and dynamic conformations in the NMR structure of the designed protein. The P-loop motifs in naturally occurring proteins typically show a cradle-like conformation to grasp the phosphate moieties of the ATP molecule (Fig. 1a). We have recently described the characteristic features of the backbone conformation of the P-loop motif (Fig. 1a)(13). Here, we hypothesize that the low ATPase ability is attributed to the unstable conformation of the inserted P-loop motif. Based on this hypothesis, we aimed to design ATPases with the P-loop that adopts the characteristic, stable conformation, by investigating the optimal backbone blueprint for de novo design of proteins harboring the P-loop motif.

**Fig. 1.**
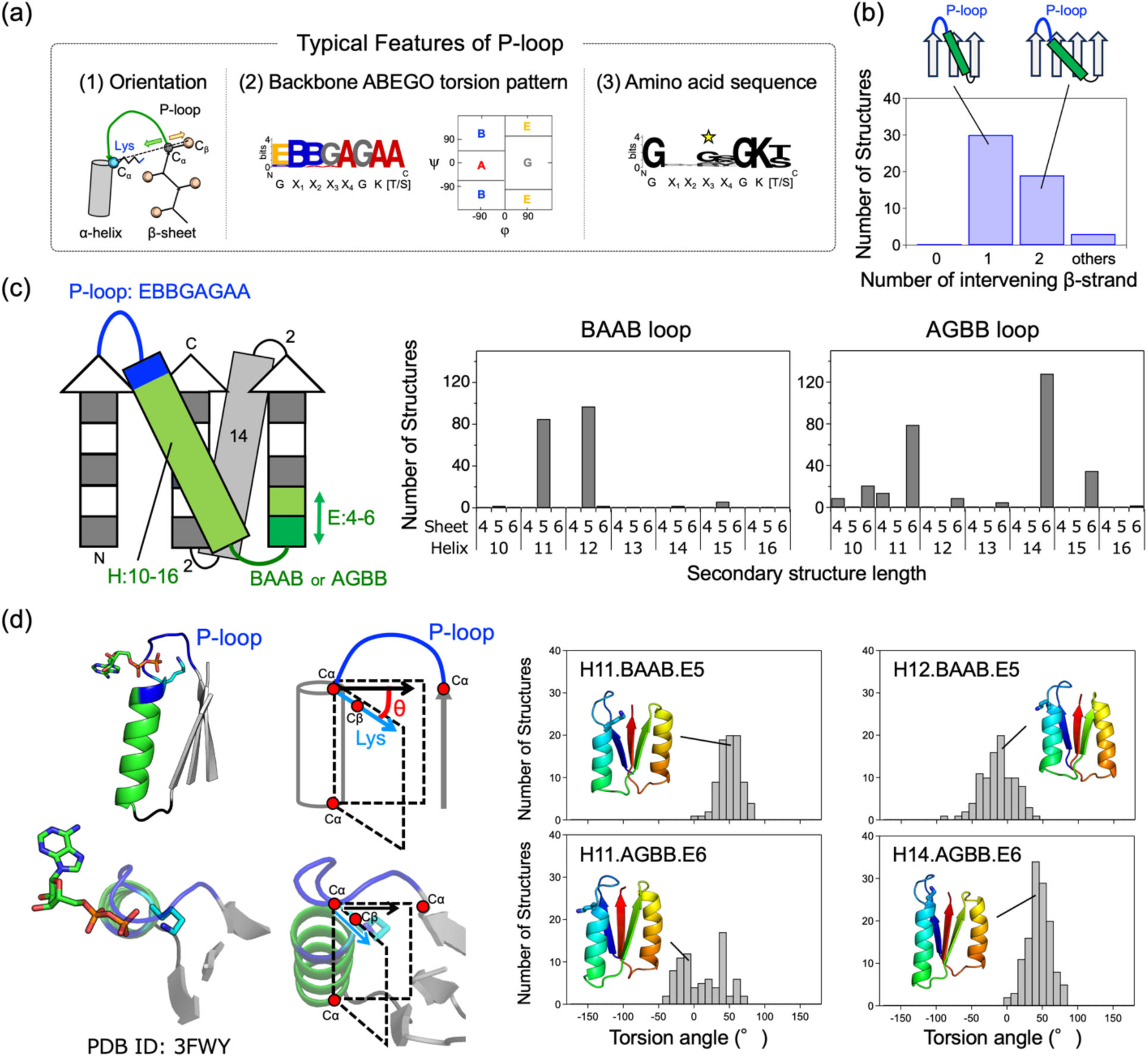
Exploration of the optimal blueprint for de novo design to harbor the P-loop motif. (a) The typical features of P-loops observed in naturally occurring proteins (13). (1) Orientation: the vector from the C_α_ atom of the last strand residue immediately before the P-loop to the C_α_ atom of the conserved Lys points away from the vector from the C_α_ atom of the same last strand residue to its C_β_ atom. (2) Backbone ABEGO torsion pattern: the backbone conformation of residues in P-loops typically show the EBBGAGAA torsion pattern (the torsion A and B are the α-helix and β-sheet regions; G and E are the positive cp regions). (3) Amino acid sequence: the P-loop is identified by the conserved sequence GX_1_X_2_X_3_X_4_GK(T/S). (b) The distribution of the number of β-strands traversed by the β-(P-loop)-α-β motifs in naturally occurring proteins. (c) Left: Explored partial blueprints to identify the optimal one to harbor the P-loop motif. Right: The distribution of the number of Rosetta folding simulation trajectories, in which the simulated protein, starting from an extended conformation, folded into the structure corresponding to each blueprint. For each blueprint, 500 independent folding simulations were carried out. (d) Left: A typical naturally occurring P-loop conformation with an ATP molecule. The conserved Lys points inward to the P-loop cradle to interact with the β-phosphate moiety of ATP. This structural feature can be captured by considering the orientation angle defined by the following two planes. One plane is defined by the Cα and Cβ atoms of the Lys residue and the Cα atom of the last residue of the helix connected to the P-loop. The other plane is defined by the Cα atom of the last β-strand residue, and the Cα atoms of the first and last helix residues. Right: The distribution of the number of structures, generated from the above Rosetta folding simulations, with the Lys showing the orientation angles between 40° and 60°.

## Results

### 2.1 Exploration of the Blueprint Optimized for Harboring the P-loop Motif

We began by seeking an optimal backbone blueprint, which involves a tertiary arrangement of secondary structures and loop connections (i.e., topology or fold), along with their lengths, to design protein structures that harbor the P-loop motif. As a first step, we attempted to build a partial blueprint for the structure around an embedded P-loop motif. The P-loop motifs are typically located at junctions from a β-strand to an α-helix in the α/β class proteins, which are characterized by a β-sheet sandwiched by α-helices, composed of repeated β-α-β motifs with β-strands mostly aligned in parallel. Additionally, in the β-(P-loop)-α-β motif, the first β-strand starts from the middle of the β-sheet, and the subsequent α-helix followed by the second β-strand traverses one or two β-strands (Fig. 1b). We studied the distribution of the number of β-strands traversed by β-(P-loop)-α-β motifs in naturally occurring proteins (Fig. 1b). We found that there are no topologies where the first and subsequent β-strands are adjacent to each other (i.e., the traversed number is zero). In previously designed proteins with the P-loop motif, this motif was embedded in β-α-β motifs where the first and second β-strands are adjacent to each other (24). Therefore, we decided to embed the β-(P-loop)-α-β motif with one intervening β-strand (Fig. 1c), considering the most common arrangement found in naturally occurring proteins. Furthermore, we previously identified typical local structural features for the P-loop motif (13): the vector from the C_α_ atom of the last strand residue immediately before the P-loop to the C_α_ atom of the conserved Lys points away from the vector from the C_α_ atom of the same last strand residue to its C_β_ atom (Fig.1a,left), and the backbone torsion pattern represented by ABEGO (32) torsion bins is EBBGAGAA (the torsion bins A and B are the α-helix and β-sheet regions; G and E are the positive phi regions) (Fig. 1a, middle). With these considerations, we explored the optimal lengths of secondary structures and the connecting loops for the α/β topology shown in Fig.1c by carrying out Rosetta fragment assembly simulations (28).

In the Rosetta folding simulations, we fixed the lengths of the first and third β-strands at five and set the loop torsion pattern of the P-loop motif to EBBGAGAA, as frequently observed in naturally occurring proteins (Fig. 1a, middle). The fourteen-residue second α-helix and the two-residue loops connected to the helix were determined by referring Table S1 and S2 in the previous work (16). Subsequently, we explored the optimal lengths of the helix connected to the P-loop motif and the second β-strand, as well as the backbone torsion patterns, represented by ABEGO torsion bins, of the loop immediately following the helix. Our aim was to identify the helix and β-strand lengths, as well as loop backbone torsion patterns, that would most frequently fold into the structures designated by the blueprints. For each blueprint, 500 independent folding simulations were carried out, resulting in the identification of four patterns of the helix length, loop ABEGO pattern, and β-strand lengths: H11.BAAB.E5, H12.BAAB.E5, H11.AGBB.E6, and H14.AGBB.E6 (where H represents a α-helix; E represents a β-strand), that consistently folded into the intended blueprint structure.

Next, we sought to identify the most optimal blueprint to harbor the P-loop among the four identified blueprints by focusing on the side-chain orientation of the conserved Lys in the P-loop motif. The P-loop motif exhibits a cradle-like conformation that encompasses the α- and β-phosphate groups of ATP, with the conserved Lys primarily interacting with the β-phosphate moiety (Fig. 1d, left). Therefore, we delved into the four identified patterns to reveal which could exhibit the optimal sidechain orientation, facilitating the binding of the conserved Lys in the P-loop motif to the β-phosphate. To this end, we examined the pattern in which the orientation of C_α_ to C_β_ vector of the Lys residue points inward to the P-loop cradle, relative to the plane defined by the C_α_ atoms of the last strand residue and the first and last helix residues. As a result, the blueprint with a 14-residue helix, a loop with the AGBB ABEGO torsion pattern, and a six-residue β-strand was identified as the most optimal for allowing the Lys residue to point inward to the P-loop cradle, facilitating the interaction with the β-phosphate moiety.

Next, we sought to construct a complete blueprint based on the identified partial blueprint, with the aim of the creation of the binding pocket for the adenine base and ribose of ATP. To achieve this, we augmented the partial blueprint by incorporating additional three β-α-β motifs (Fig. 2a). Register shifts between the fourth and fifth strands, as well as between the fifth and sixth strands, were introduced to facilitate the twisting of the β-sheet, allowing the C-terminal helix to interact with the N-terminal and with the adenine ring of ATP. The lengths of secondary structures and the connecting loops were selected by referencing Table S1 and S2 in the previous work (16), except for the last helix and its connected loop. The length of the last helix was chosen to sufficiently cover the β-sheet. In our previous de novo designed proteins, loops connecting a β-strand to an α-helix were either two or three residues, depending on the helix’s orientation relative to the C_α_ to C_β_ vector of the last strand residue (8). However, for this design, we opted for a loop length of four residues to provide enhanced conformational freedom to search for conformations in which the loop and following helix can interact with the ATP molecule.

**Fig. 2.**
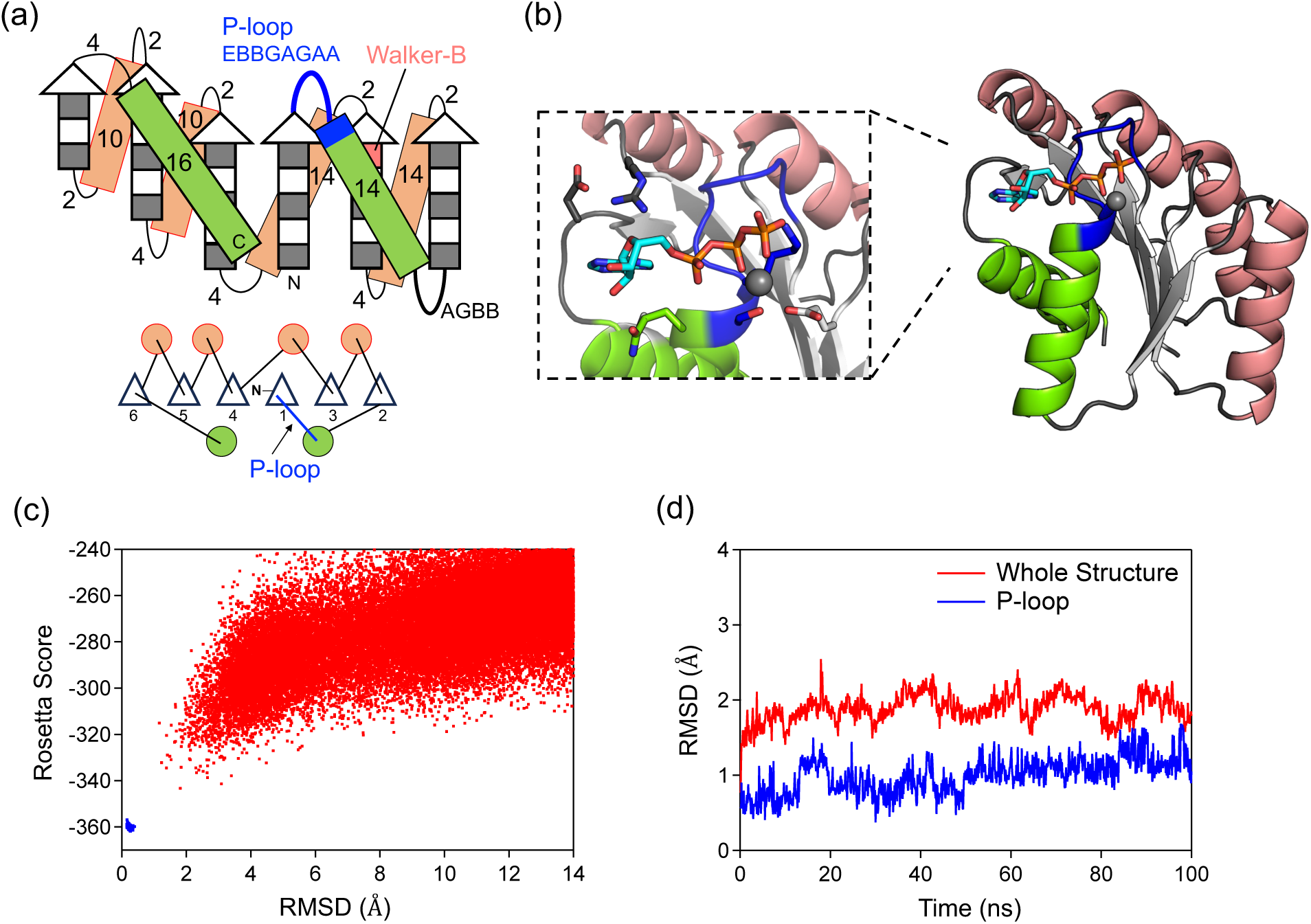
De novo designed protein with the P-loop motif. (a) The identified optimal backbone blueprint to harbor the P-loop motif. Strand lengths are represented by filled and empty boxes that indicate pleats coming out and going into the page, respectively. Helices are represented by rectangle boxes. Letter strings next to the loops or on the rectangle boxes indicate their lengths. (b) The designed protein structure based on the optimal blueprint. This structure harboring the P-loop motif also contains a binding pocket for the base and ribose moieties, allowing it to encompass an entire ATP molecule. (c) The funnel-shaped energy landscape of the designed protein, PL2×4_2, explored by Rosetta folding simulations. Lowest-energy structures obtained by 50,000 and 5,000 independent Monte Carlo structure prediction trajectories starting from an extended chain (red) and from the design model (blue) are shown. The x and y axes indicate the Cα root mean squared deviation (RMSD) from the design model and the Rosetta all-atom energy, respectively. (d) Conformational fluctuations of PL2×4_2 during MD simulations for the whole structure (red) and the structure of P-loop motif (blue). Cα RMSD values from the initial structure during the simulations are shown.

### 2.2 Design of ATPase based on the Identified Optimal Blueprint

Finally, we built protein structures based on the complete backbone blueprint (Fig. 2). We carried out Rosetta fragment assembly simulations, which generated approximately 100 backbone structures harboring the P-loop motif. For each generated structure, amino acid sequences and their side-chain conformations were built using RosettaDesign (15) to stabilize the entire generated backbone structure as well as to make favorable interactions with the ATP molecule, using multiple ATP conformations and a set of distance constraints used in our previous research (13). We also attempted to place the Walker-B motif (i.e., Asp or Glu), which chelates the magnesium ion with the Thr or Ser residues of the P-loop motif, at the second-to-last residue of the third β-strand.

The designed structures were filtered based on the binding scores (Roestta ddG score < -12.0, DSasa score >0.4 and <0.7), the side chain packing quality (RosettaHole score <2.0, packstat > 0.6) (27), the compatibility of sequence and structure (8). One of the designed structures is shown in Fig 2b. Subsequently, we selected designs with funnel-shaped energy landscapes, explored through Rosetta ab initio structure prediction simulations (23) for the designed structures in the absence of an ATP molecule (Fig. 2c). The stability of both the backbone structure and the P-loop motif was further evaluated by investigating the structural stability during a 100ns MD simulation (Fig. 2d). Finally, three designs that satisfy all of the above criteria were selected for experimental evaluation.

### 2.3 Experimental Characterization of the designed ATPases

We obtained synthetic genes encoding the three designs (Supplementary Table 2). None of the designs showed similarity to known proteins, with a BLAST E-value < 0.001. The proteins were recombinantly expressed in *Escherichia coli* and purified using Ni^2+^-affinity (Ni-NTA) chromatography. The purified proteins were characterized using circular dichroism (CD) spectroscopy, size-exclusion chromatography combined with multi-angle light scattering (SEC-MALS), and ^1^H-^15^N heteronuclear single quantum coherence (HSQC) NMR spectroscopy. All designed proteins were found to be expressed and highly soluble, and exhibited CD spectra typical of αβ-proteins from 20 °C to 98 °C.

Two of the three designs PL2×4_1 and PL2×4_2 were found to be a monomeric state in solution by SEC-MALS. PL2×4_2, which was more soluble than PL2×4_1, displayed well-dispersed sharp NMR peaks, indicating the folding of the design into rigid tertiary conformations. Accordingly, we attempted to crystalize the design, PL2×4_2, and determined the structure at 2.4 Å resolution (PDB 9JIX). The crystallographic data reveal two molecules in an asymmetric unit. Each monomer shows a slightly different structural state, referred to as Crystal1 and Crystal2 (Fig. 3). Comparison of the designed model with these crystal structures demonstrates good agreement between the designed and experimental structures, except for the fifth helix and sixth strand and the loops connected to the helix and strand (Fig. 3d). Moreover, the P-loop motif in the structure exhibited the typical features of the P-loop motif (Fig. 3e). Extra densities near the P-loop motif suggested the possible presence of a bound sulfate, because the crystallization condition contained ammonium sulfate (Fig. 3f).

**Fig. 3.**
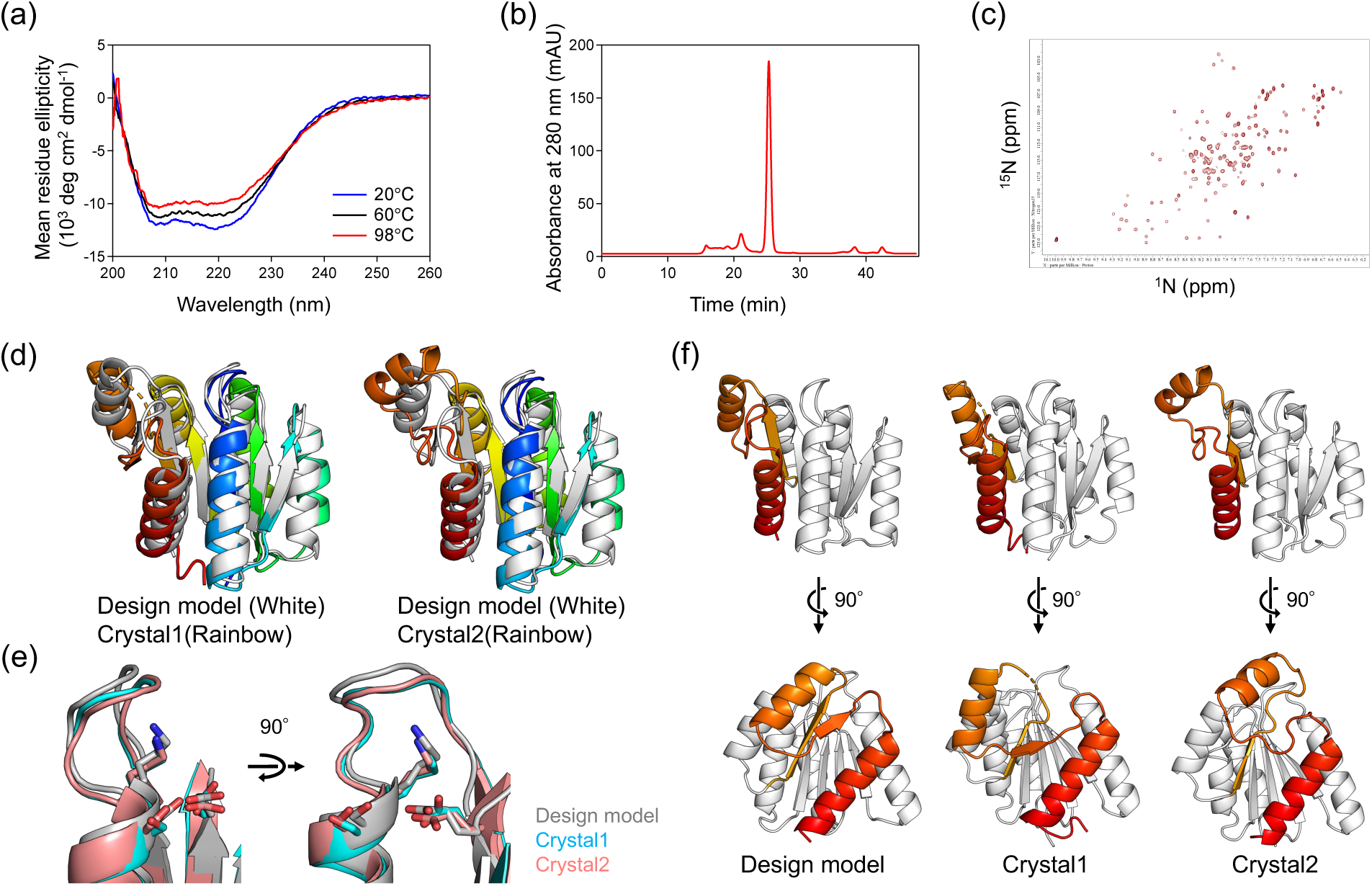
Experimental characterization of the designed protein, PL2×4_2. (a) Far-ultraviolet CD spectra at temperatures, 20 °C, 60 °C, and 98 °C. (b) UV signal from the SEC-MALS measurement. (c) Two-dimensional ^1^H-^15^N HSQC NMR spectra at 25 °C and 600 MHz. (d+f) Comparison of the design and its crystal structures. The crystal structures reveal two molecules (Crystal 1 and 2) in in an asymmetric unit, demonstrating different structural states. The Cα RMSD values between the design model and Crystal 1, and the design model and Crystal2, are 1.8 and 1.7 Å, respectively. (e) Close-up view of the design and crystal structures focusing on the P-loop motif.

We also predicted the folded structure of the designed protein PL2×4_2 using AlphaFold2 (6). Interestingly, the predicted structure exhibited a different topology from the designed structure, which was confirmed to be correct by X-ray crystallography, specifically in the order of β-strands (Fig. 4). No homologs are present in naturally occurring proteins for our designed protein. Given that AlphaFold is primarily trained on naturally occurring proteins, the absence of homologs posed a challenge in predicting the folded structure.

**Fig. 4.**
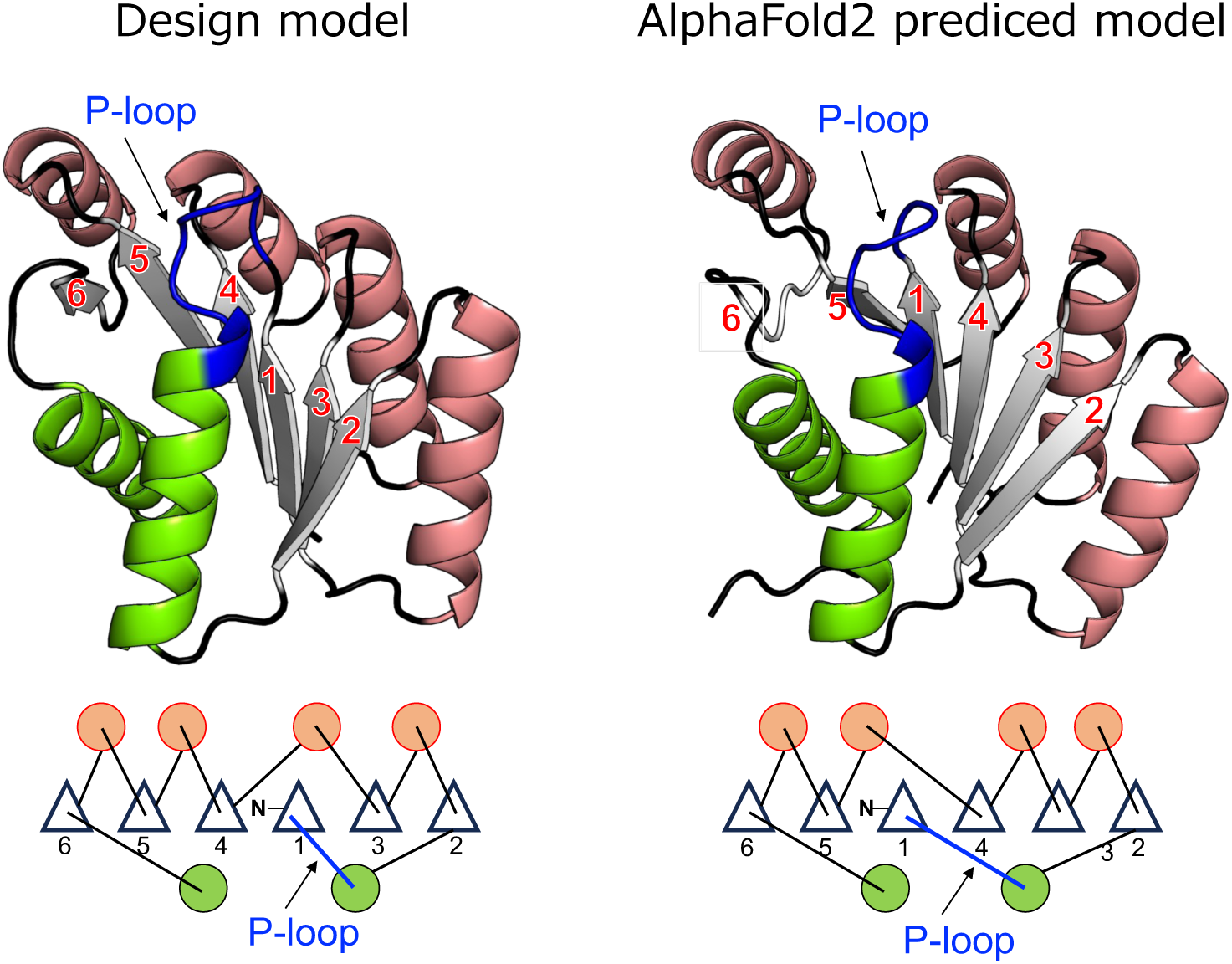
Comparison of the design model (left) with the AlphaFold2 predicted model (right) The structure and its topology of our design model (left) and those of the AlphaFold2 predicted model (right) are shown. The order of β-strands in the predicted model is different from that of the design model: the β-(P-loop)-α-β motifs in the design and predicted models traverse one and two intervening β-strands, respectively.

Next, the binding affinity of PL2×4_2 for ADP was measured using the fluorescence polarization method with the fluorescent-labeled ADP (Mant-ADP) (Fig. 5a). Clear binding signals for the design were detected, indicating that ADP likely binds to the design. Moreover, the mutant for the conserved Lys residue, K14Q, in the P-loop motif exhibited a lower signal than the original design, suggesting that the design binds an ADP molecule in proximity to the P-loop motif. However, the determination of the exact *Kd* values was challenging due to the expected low affinity, approximately a few mM; it was difficult to prepare protein samples at sufficiently high concentration. The approximate *Kd* values from the current signals were estimated to be 1.46 ± 0.30 mM for the design and 1.88 ± 0.51 mM for the K14Q mutant. For reference, the *Kd* value of F_1_-ATPase from thermophilic *Bacillus* PS3 is 19 ± 1 μM (34), and the median *K_M_* value of naturally occurring enzymes is about 100μM (18).

**Fig. 5.**
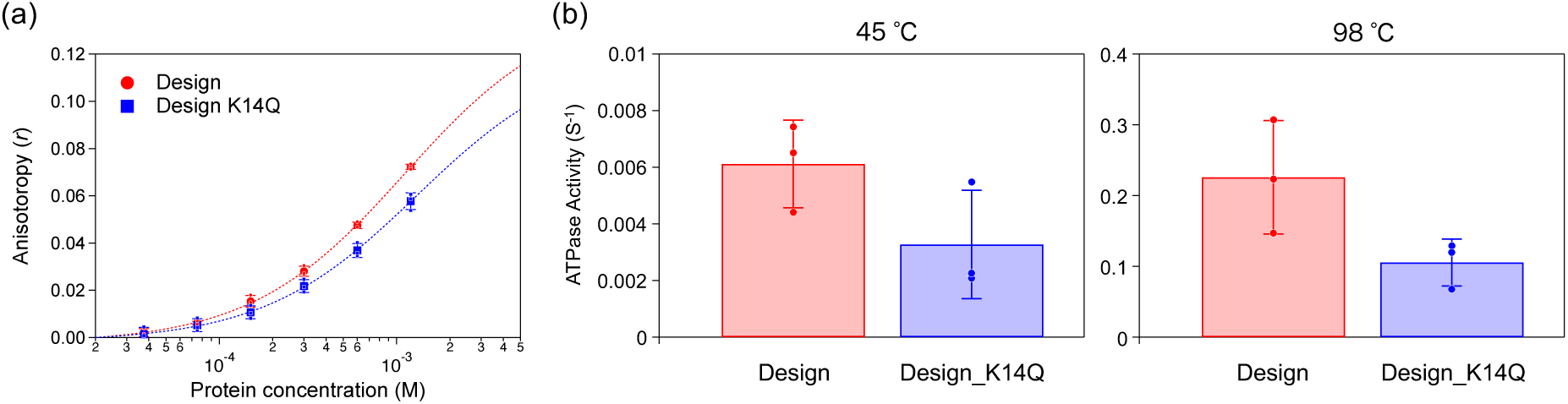
ATP binding and hydrolysis activities of the designed protein, PL2×4_2. (a) ATP binding assays measured by fluorescence polarization for PL2×4_2 (red) and its K14Q mutant (blue). (b) ATP hydrolysis assays conducted using the ATPase/GTPase Activity Assay Kit (Sigma-Aldrich Co. LLC, MAK113) for PL2×4_2 (red) and the K14Q mutant (blue) at temperatures 45 °C and 98 °C.

Finally, we measured ATP hydrolysis ability of the design by detecting the product, phosphate molecules, using the malachite green reagent (Fig. 5b). The design exhibited weak ATP hydrolysis activity; the rate of ATP hydrolysis is approximately 0.0061 ± 0.0015 min^-1^ at 45 °C. The P-loop structure in our design showcased the typical features observed in naturally occurring proteins, as observed in the crystal structure (Fig. 3d). Moreover, the decreased binding affinity in the K14Q mutant supports that the P-loop motif in the design adopts the typical features. Therefore, the designed P-loop was expected to enhance the ATP hydrolysis capability. However, the activity was comparable to those of previously designed proteins (24) (31): the ATP hydrolysis rate of the ATPase designed by Romero et al. is 0.0058 ± 0.0012 min^-1^ at 45 °C (24) and the *k_cat_* value of the ATPase designed by Wang et al. is 0.00058 ± 0.00005 min^-1^ at room temperature (31). To further investigate, we tested the activity of the design at a higher temperature (98 °C). At this temperature, the hydrolysis activity drastically increased to approximately 0.23 ± 0.08 min^-1^. This increase in activity can be attributed to the high-temperature stability of the design, although its activity levels still fall short compared to those of naturally occurring enzymes. For reference, the V_max_ value of F_1_-ATPase from thermophilic *Bacillus* PS3 is 247 ± 9 s^-1^ at 23 °C (34), and the median *k_cat_* values of naturally occurring enzymes is about 1.0 x 10^2^ s^-1^(18).

## Discussion

We aimed to design an ATPase with the P-loop motif by investigating the optimal backbone blueprint to harbor the P-loop motif in designed structures. To achieve this, we conducted a statistical analysis of naturally occurring proteins with the P-loop motif and performed Rosetta folding simulations. Based on the identified backbone blueprint, we designed three proteins, two of which successfully folded into monomeric structures with high thermal stability. The crystal structure of one of these designs, PL2×4_2, was determined and found to be nearly identical to the design model, both in the overall structure and the P-loop motif. As expected, PL2×4_2 demonstrated ADP binding and ATP hydrolysis abilities. However, these abilities are lower than those of typical naturally occurring ATPases.

We examined the reasons for this reduced functionality. For this purpose, we conducted MD simulations for the designed protein complexed with an ATP molecule. The simulations suggested that the low binding and hydrolysis activities might be due to a suboptimal binding pocket for the adenine and ribose moieties of ATP, as these moieties were detached from the binding pocket during the simulations (Fig. 6). Furthermore, we found that the loop (the C-terminal loop connected the sixth strand) adjacent to the binding pocket exhibited unsteady dynamic motion. This motion could contribute to the weak binding of the pocket against the adenine and ribose moieties. Consistent with the simulations, each monomer in our crystal structure revealed different loop conformations, suggesting that the loop’s flexibility might indeed contribute to the suboptimal binding pocket. While our identified blueprint was optimal for building structures that harbor the P-loop motif, it may not be ideal for forming the binding pocket. Another possibility is the absence of the Arg-finger, which is known to be a crucial component of the ATP hydrolysis module(10) that works with the P-loop and Walker-B motifs. Additionally, protein dynamics that facilitate the catalytic activity needs to be considered (12). The blueprint we developed in this work were for creating ideal proteins completely optimized for folding ability, but longer loops or local frustrations encoding functional dynamics can be important to achieve high ATPase activity. Future investigations should focus on blueprints (and amino acid sequences) that include these features.

**Fig. 6.**
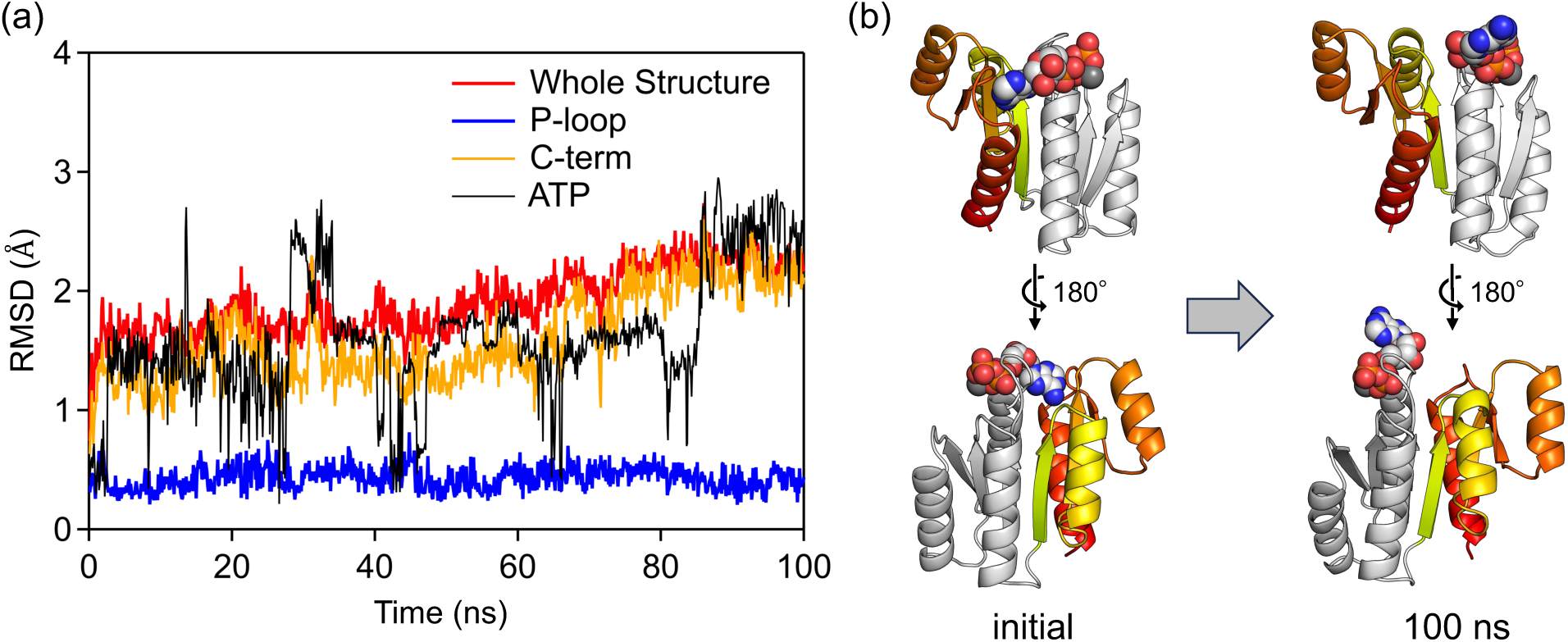
The created binding pocket for adenine of ATP may not be optimal. (a) MD simulations of the designed PL2×4_2 complex with an ATP molecule. The RMSD values for the entire structure (red), the P-loop motif structure (blue), the C-terminal structure (orange) and ATP molecule (black) are shown. While the P-loop motif maintains its initial conformation during the simulation, the conformations of the C-terminal and the ATP molecule change. (b) The initial and final structures from the MD trajectory are depicted. The adenine ring of the simulated ATP molecule (sphere) was released from the designed binding pocket. The pocket was collapsed by change of the C-terminal structure (colored region in the protein structure).

## Materials and methods

### Backbone building and sequence design protocol

The backbone structures for the blueprint were built by Rosetta sequence-independent folding simulations using coarse-grained model structures, in which each residue is represented by main chain atoms (N, H, CA, C and O) and a side chain pseudo atom (23). The following terms were included in the Rosetta potential function used in the simulations with no sequence-dependent score terms; steric repulsion (vdw = 1.0), overall compaction (rg =1.0), and hydrogen bonds (hbond_sr_bb = 1.0, hbond_lr_bb = 1.0). The steric residues of Val was used for that of the side chain pseudo atom. An ATP molecule for each generated main-chain structures was placed near the P-loop motif using the Rosetta PredesignPerturb protocol, which randomly perturb a ligand in a protein active site, with the distance constraints used in our previous research (13). Sequence design were performed by RosettaDesign calculation (15) using the flexible-backbone design (FlxbbDesign) protocol with the full-atom Talaris2014 scouring function (22). The definition of the environment for each residue position, layer, and the designated amino-acids for each layer are almost same with the previous paper (8), except for the point that we excluded large aromatic residues, F and W, form the boundary layer in this paper. We also added constraints for the strand pairs to keep the backbone structure. The parameters for ATP-Mg^2+^ were determined using those for the corresponding atom types defined in Rosetta. A set of various ATP conformations was generated by BCL software(14).

### Molecular dynamics (MD) simulations

The AMBER14 software suite (4) was used for all MD simulations. The design models were used as the initial structures, of which hydrogen atoms were added by the LEaP module of AMBER14. The simulation system contains a designed monomer with ATP placed in a water box of approximately 65 Å × 75 Å × 65 Å. To neutralize the system, 1 or 2 sodium ions or 2 chloride ions were put in the box. AMBER ff99SB (first screening) or ff14SB (evaluation of ATP binding) sets and TIP3P were utilized for the protein and water molecules, respectively. Parameters for ATP molecule were adopted from a reference paper(20). Long range electrostatic interactions were treated by the particle mesh Ewald (PME) method. Non-bonded interactions were cut off at 10 Å. After carrying out a short minimization to remove artificial repulsions in the initial structure, 100 ns MD simulations in a constant-NPT (300K, 1atm) ensemble were performed after the 100 ps heating stage with NVT ensemble (the time step is 2.0 fs and hydrogen atoms were constrained with SHAKE procedure). At the heating step, the temperature was raised gradually from 0 K to 300 K with the weak restraints (10 kcal/mol/A^2^) to the atoms of the designed protein.

### Plasmid construction, expression, and purification of designed proteins

The genes encoding the designed sequences (Supplementary Table 2) were synthesized and cloned into pET21b vectors (Eurofins Genomics). *E. Coli* BL21* (DE3) competent cells were transformed with the plasmid and cultured at 37 °C for 6 hour followed by overnight incubation at 18 °C in autoinduction media(29). The expressed proteins were purified using Ni-NTA column and then dialyzed against Tris buffer (10 mM Tirs-HCl (pH 8.0), 100 mM NaCl and 5 mM MgCl_2_). The purified protein samples were stored at -80 °C. For ATP-binding and ATPase activity assay, the samples were concentrated with a Vivaspin20 5,000 molecular weight cut-off (MWCO) (Cytiva) and then passed through a Superdex 75 Increase column (Cytiva) equilibrated with Tris buffer (pH 7.4). The purified samples were stored at -80 °C.

### Plasmid Construction for the design mutants

Mutations were introduced by the Quick Change Multi Site-Directed Mutagenesis Kit from Agilent Technologies. The expression and purification for the mutants were carried out by the same protocol as for the original design. The DNA sequence was confirmed by DNA sequencing analysis performed by Fasmac.

### Circular dichroism (CD) measurement

All CD measurements were performed using a JASCO J-1500 CD spectrometer, using in a 1mm path length cuvette. Far-UV CD spectra for the designed proteins were measured in the wavelength range from 260 to 200 nm. The measurements were conducted at various temperatures ranging from 25 °C to 98 °C, for protein samples at concentrations of 10-50 μM in Tris buffer (pH 8.0).

### SEC-MALS

SEC-MALS experiments were performed using a miniDAWN TREOS static light scattering detection system from Wyatt Technology Corporation, combined with a HPLC system (1260 Infinity LC; Agilent Technologies) with a Superdex 75 increase 10/300 GL column from Cytiva. After equilibration of the column with PBS buffer (pH 7.4), the volume 100 μL of protein samples (300 ∼ 500 μM) in PBS buffer (pH 7.4), which were purified beforehand by Ni-affinity columns, was injected. Protein concentrations were calculated from the absorbance at 280 nm detected by the HPLC system. Static light scattering data were collected at three different angles, 43.6°, 90.0°, and 136.4°, using a 659 nm laser. These data were analyzed by the ASTRA software (version 6.1.2, Wyatt Technology Corporation) with a change in the refractive index with concentration, a *dn/dc* value, 0.185 mL/g to estimate the molecular weights of the dominant peaks.

### Expression and purification of protein sample for NMR measurements

The designed proteins were expressed in *E. Coli* BL21* (DE3) cells as uniformly ^15^N-labeled proteins. These ^15^N-labeled proteins were expressed by using MJ9 minimal media, which contains ^15^N ammonium sulfate as the sole nitrogen source and ^12^C glucose as the sole carbon source. The expressed proteins with a 6xHis tag were purified using Ni-NTA column and then dialyzed to Tris buffer (10 mM Tris-HCl (pH 8.0), 100 mM NaCl and 5 mM MgCl_2_). The samples were concentrated with a Vivaspin20 5,000 molecular weight cut-off (MWCO) (Cytiva) and then passed through a Superdex 75 Increase column (Cytiva) equilibrated with PBS buffer (pH 7.4). The purified samples were stored at -80 °C.

### 1H-15N HSQC NMR spectroscopy

To confirm the core packing of the designed proteins, we measured 2D ^1^H-^15^N HSQC spectra for the most promising design. The spectra were collected for 400-500 μM samples in 90% ^1^H_2_O/10% ^2^H_2_O PBS buffer (pH 7.4) at 25 °C on a JEOL JNM-ECA 600 MHz spectrometer and were processed and analyzed using JEOL Delta NMR software.

### Expression and purification of protein sample for crystallization

The protein sample for crystallization was prepared by removing the His-tag from the N-terminal of the design protein. The gene encoding the designed PL2×4_2 sequence was cloned into the pET21b vector and then cloned into the pET15b vector. A TEV protease cleavage site and a Gly residue were inserted between the gene encoding the designed protein and the His-tag in the pET15b vector. Using this plasmid, the designed protein was expressed and subsequently purified via a Ni-NTA column. The eluted sample was mixed with TEV protease in a 10:1 molar ratio and dialyzed against a 20 mM Tris-HCl buffer (pH 8.0). This dialyzed samples were applied to a Ni-NTA column, and the flow through was collected. Next, the sample was loaded onto a HiTrap Q HP column (Cytiva) equilibrated with 20 mM Tris-HCl buffer (pH 8.0), and then eluted with a linear gradient of Tris buffer with 0-1000 mM NaCl in 20min at a flow rate of 1.0 ml min^-1^. The concentrated sample with a Vivaspin 20 (5,000 MWCO) (Sartorius) was loaded onto a Superdex 75 Increase 10/300 GL column (Cytiva) equilibrated with Tris buffer without MgCl_2_ (10mM Tris-HCl (pH 8.0) and 100 mM NaCl) at a flow rate 0.5 ml min^-1^. The purified sample was concentrated using a Vivaspin 500 (5,000 MWCO).

### Crystallization, data collection and structure determination

The sitting drop vapor diffusion method was used for crystallization. Crystals of PL2×4_2 were obtained by mixing a 2.0 μL protein solution (37.7 mg/mL protein in Tris buffer) with 2.0 μL of the reservoir solution (0.1 M Sodium Acetate (pH 4.5) and 2.0 M Ammonium Sulfate), which was equilibrated against 500 µl of reservoir solution. The crystals appeared in 1-2 weeks at a temperature of 293K. The reservoir solution with 40% (v/v) concentration of glycerol was gradually added to the drop containing crystals up to 20% (v/v) concentration.

The crystals were mounted on cryo-loops (Hampton Research), flash-cooled, and stored in liquid nitrogen for preservation. X-ray diffraction data were collected using a wavelength of 1.1 Å on BL-1A of the Photon Factory (PF) using a single crystal at the cryogenic temperature of 100K. The collected data were processed by XDS (7). The structure was determined by the molecular replacement method with Phaser (19), using the designed model as an initial search template. The initial model was iteratively refined with PHENIX (1) and REFMAC5 from the CCP4 Suite (21) and manually corrected using COOT (5). The figures in the manuscript were generated by PyMOL (26). The crystallographic and refinement statistics are summarized in Supplementary Table 1.

The crystal structures have been deposited in the wwPDB as PDB 9JIX.

### Fluorescence polarization measurement for evaluating ADP-binding affinity

Fluorescence polarization-based affinity measurements for the designed proteins were performed using 1 μM fluorescent-labeled ADP, Mant-ADP. We observed changes in fluorescence anisotropy (*r*) of the fluorescent-labeled ADP mixed with the protein samples at varying concentrations. The mixtures were prepared in Greiner black flat bottom 96-well plates and equilibrated 1 hr at room temperature. Measurements were carried out on a Spark 10M (TECAN), using 360 nm excitation and 464 nm emission, each with a 35 nm bandwidth. All measurements were conducted in Tris buffer.

Equilibrium dissociation constants (*K_d_*) were determined by fitting the observed anisotropy data, which was obtained 20 measurements over a 10 minute period for each protein concentration, to Equation 1. In this equation, *A* is experimentally measured anisotropy, *A_f_* is anisotropy of the free ligand, *A_b_* is the anisotropy of the fully bound ligand, [*L*]*_T_* is the total ligand concentration, and [*R*]*_T_* is the total protein concentration. The *K_d_* values were determined by averaging the values from three independent measurements.

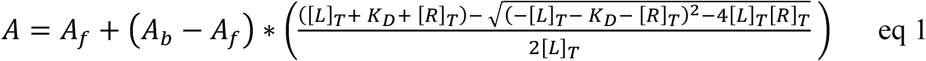

### ATPase activity measurements

ATPase activities for the designed proteins were measured by the ATPase/GTPase Activity Assay Kit (Sigma-Aldrich Co. LLC, MAK113). Protein samples at concentrations of 25 μM (for reactions at 45 °C) or 5 μM (for reactions at 98 °C) were incubated with 1 mM ATP in 40 μL Assay Buffer (40 mM Tris, 80 mM NaCl, 8 mM MgAc_2_, 1mM EDTA, pH 7.5) in 96 well plates (Greiner, 665801) for 30 min at 45 °C or 5 min at 98 °C, respectively. As a reference, solutions in the absence of protein samples were also incubated at the same conditions. After incubation, the samples were immediately put on ice, 200 μL of malachite green reagent was added to each well, and the mixtures were incubated for 30 min at room temperature. The absorbance of the incubated solutions was measured between 350-850 nm using a Spark 10M (TECAN). The product (phosphate) concentration was estimated from the relative absorbance at 620 nm, using the reference solution for comparison against standard buffers at known several phosphate concentrations. ATPase activities for the designed proteins were calculated from the estimated product concentrations.

## Supplementary material description

Supplementary Information is available for this paper. The designed model structure will be provided with the accepted paper.

## Acknowledgements

We thank K. Nakamura and H. Yamada at SOKENDAI and D. Baker at University of Washington for discussion regarding the design of P-loop containing proteins; N. Suzuki at IMS for her assistance in sample preparation; R. Koga at Osaka University for general discussion. We thank the Research Center for Computational Science (RCCS), Okazaki, Japan for providing its computational resources; the staff at the Photon Factory (PF) at KEK for assistance in synchrotron experiments and data collection for crystallographic analyses under the approval of PF program advisory committee (Proposal No. 2017G141); and for the support of the Basis for Supporting Innovative Drug Discovery and Life Science Research (BINDS) from AMED under Grant Number JP21am0101071 (supporting number BINDS0532). This work was supported by a Grant-in-Aid for Scientific Research on Innovative Areas “Molecular Engine” (JSPS KAKENHI Grant Number 18H05420 to T. K. and N. K.), and by the Japan Science and Technology Agency (JST) Precursory Research for Embryonic Science and Technology (PRESTO, Grant Number JPMJPR20E6 to T. K.).

