## Supplementary Information for "De novo design of ATPase based on the blueprint optimized for harboring the P-loop motif"

### Table of Contents

Supplementary Figure 1: Characterization of all designs

Supplementary Table1: Data collection and refinement statistics of crystal structure

Supplementary Table2: Amino acid sequences of designed proteins

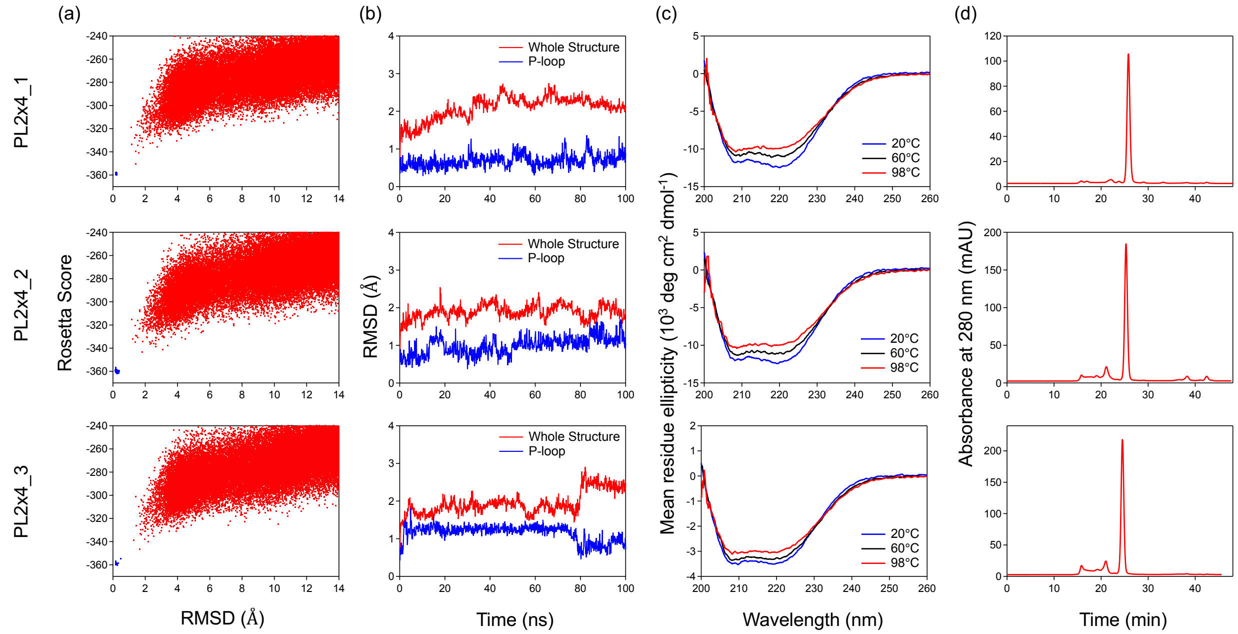

**Supplementary Figure 1. Characterization of all designs**

(a) Energy landscapes from Rosetta ab initio structure prediction simulations. (b) Cα root mean square deviation (RMSD) values during MD simulations for designs without ATP. (c) Far-ultraviolet circular dichroism (CD) spectra at various temperatures. (d) UV signals from size-exclusion chromatography combined with multi-angle light scattering (SEC-MALS).

**Supplementary Table 1: Data collection and refinement statistics of crystal structure**

| | **PL2x4_2 PDB: 9JIX** |
|---|---|
| **Data collection** | |
| **Space group** | $P3_121$ |
| **Cell dimensions** | |
| ***a, b, c*** **(Å)** | 77.66, 77.66, 101.54 |
| **α, β, γ (°)** | 90.0, 90.0, 120.0 |
| **Wavelength** | 1.070 |
| **Resolution (Å)** | 40.52 – 2.29 (2.37 – 2.29) |
| $R_{merge}$ | 0.059 (1.012) |
| ***I/σI*** | 18.3 (2.3) |
| $C/C_{1/2}$ | 1.000 (0.785) |
| **Completeness (%)** | 100.0 (100.0) |
| **Redundancy** | 9.8 (10.4) |
| **Refinement** | |
| **Resolution (Å)** | 40.52 - 2.29 |
| **No. reflections** | 16412 |
| $R_{work}/R_{free}$ | 0.237/0.260 |
| **No. atoms** | |
| **Protein** | 2308 |
| **Ligand/ion** | 31 |
| **Water** | 19 |
| **B-factors** | |
| **Protein** | 79.4 |
| **Ligand/ion** | 90.0 |
| **Water** | 57.8 |
| **R.m.s. deviations** | |
| **Bond length (Å)** | 0.011 |
| **Bond angles (°)** | 1.602 |

A single crystal was used to obtain data set. Values in parentheses are for highest-resolution shell.

**Supplementary Table 1:Amino acid sequences of designed proteins**

Computationally designed amino acid sequences are shown in uppercase and amino acid residues added to allow expression, purification, concentration measurement, cleavage sites of restriction enzymes and the spacer between the designed sequence and the C-terminal His-tag are shown in lowercase.

| Name | Amino acid sequence |
|---|---|
| **PL2x4_1** | mgAIVILVVGPPGSGKSQLIEAIERLARKQGQPVVTTSVTSEDEAKKVLRHLLKRDPNAIVVIEIKSPSIAERVAEEVLRQDPTAVLVVVVSSPDQARKLREQLPNVIVVVLIRDPEKLKEAKKEGTQVLSGNGNPEEAAKIIAQLIKDQAgswslehhhhhh |
| **PL2x4_2** | mgAIVILVVGPPGSGKSQLIKAIEKLAREQGQPVVTTSVTSEDEAKKVLEELLKKDPNAIVVIEIKNPRIAERVAKRVLEEDPTAVLVVVVSSPEVARELRENLPNVIVVVLIRDPEKLKEAKKQGTQVLSGDGNPEEAAKQIAQLIKDQAgswslehhhhhh |
| **PL2x4_3** | mgAIVILVVGPPGSGKSQLIKAIEKLAREQGQPVITTSVTSEDEAKEELERLLKKDPNAIVVIEIKSSRIAERVAKRVWEEDPTAVLVVVVSSPEDARELRENLPDVIVVVLIRDPEKLKEAKKEGTQVLSGNGNPEEAAKIIAQLIKDQAgslehhhhhh |
